## supplementary figure 1-6 for "A cleaved METTL3 potentiates the METTL3-WTAP interaction and breast cancer progression"

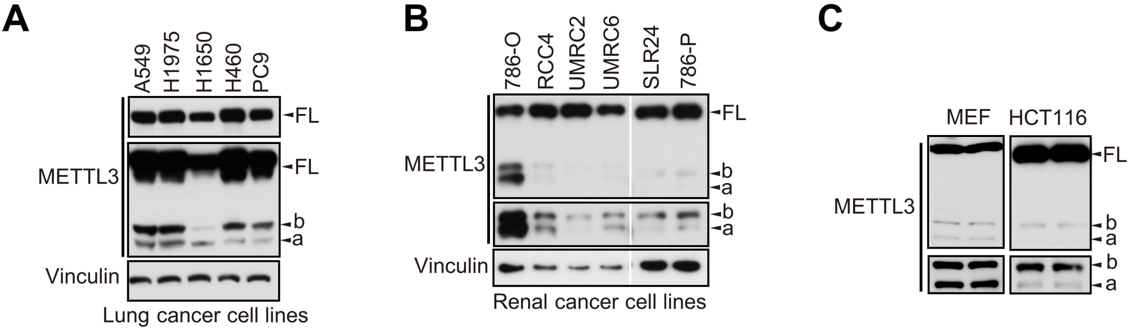

### Supplementary Figure 2

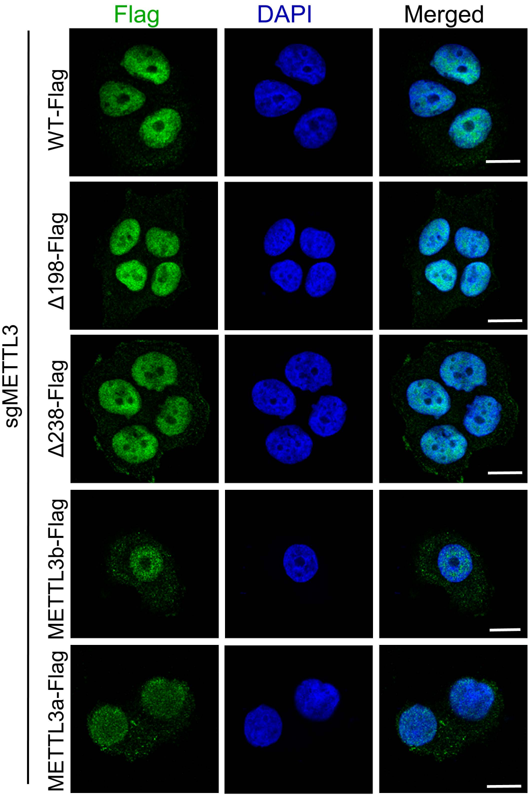

### Supplementary Figure 3

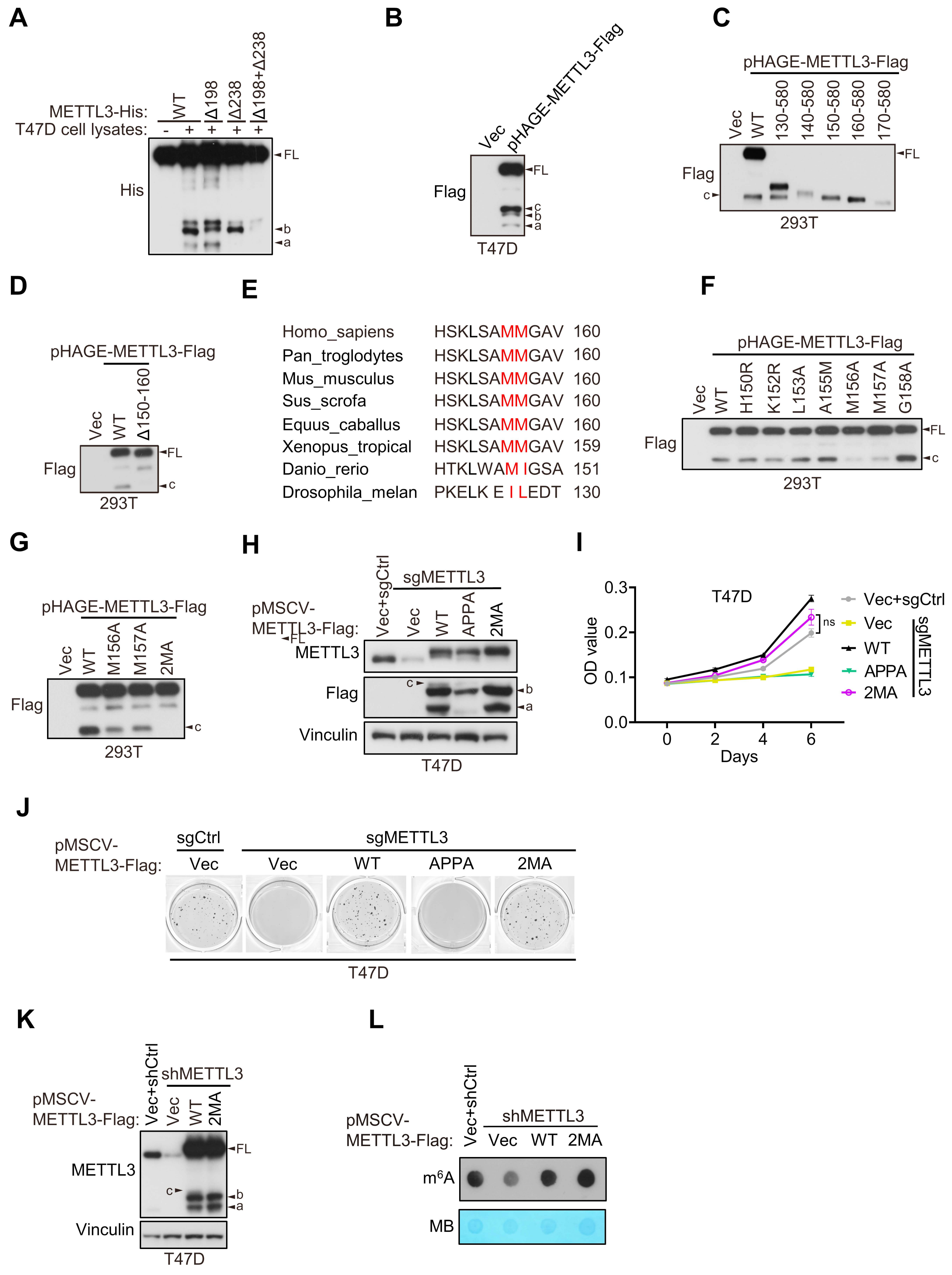

### Supplementary Figure 4

A

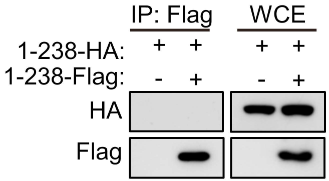


B

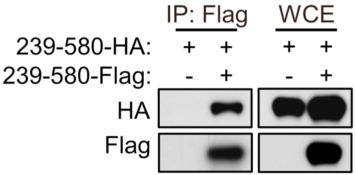


C

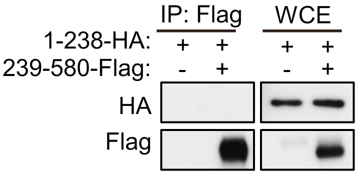


D

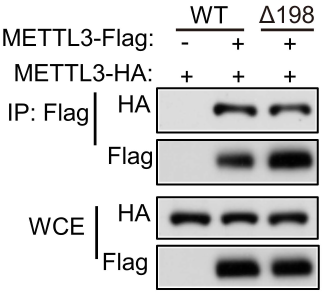

### Supplementary Figure 5

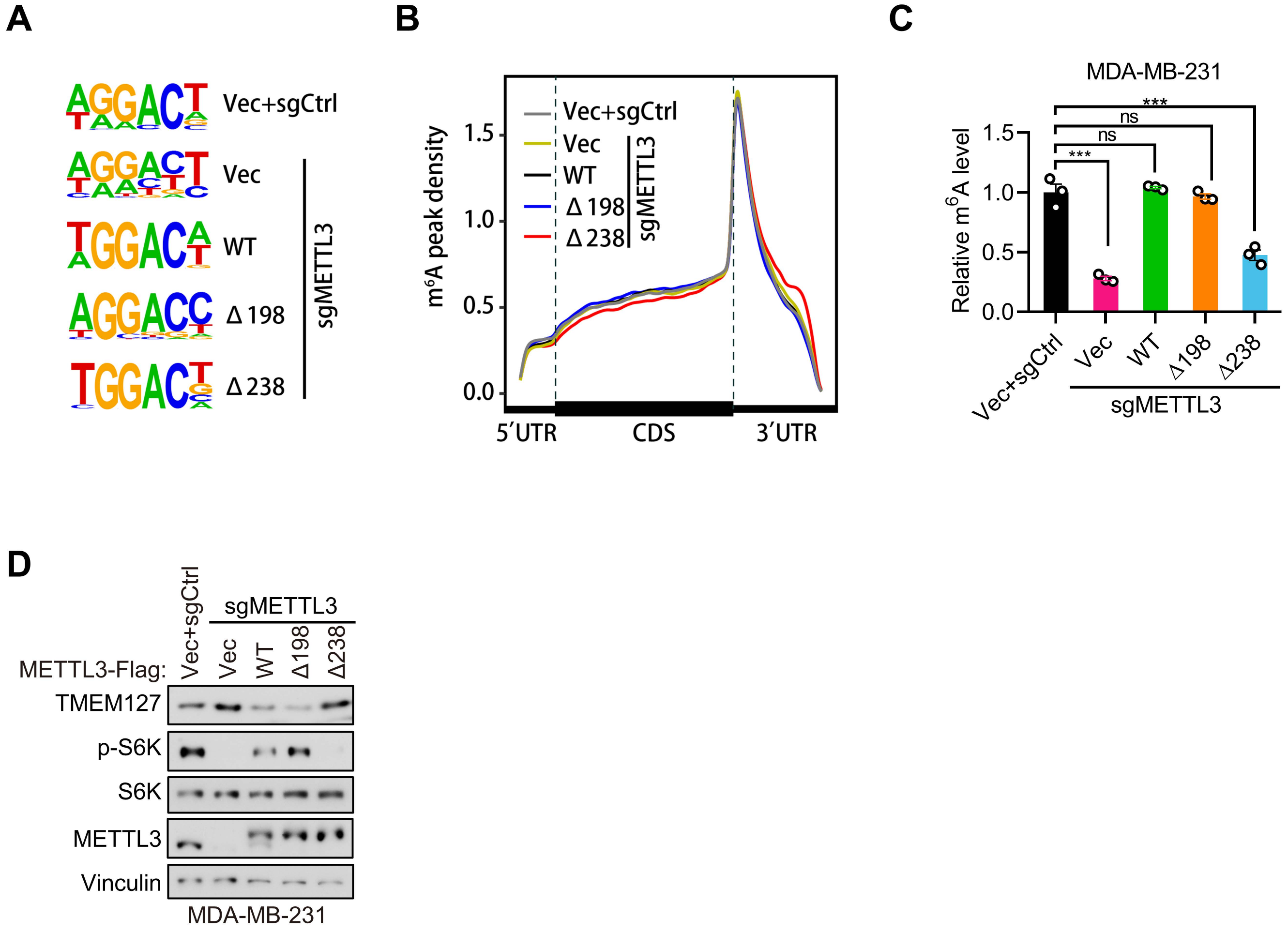

### Supplementary Figure 6

**A**

| cat. Number | Name | Target |
|---|---|---|
| **S3692** | N-Ethylmaleimide (NEM) | Cysteine Protease |
| **S3025** | PMSF | irreversible serine/cysteine protease |
| **S8102** | Z-VAD(OH)-FMK | irreversible pan-caspase inhibitor |
| **P2550** | WAY-327947 | caspase |
| **S8565** | Omarigliptin (MK-3102) | DPP-4 |
| **S8455** | Talabostat (Val-boroPro, PT-100) | dipeptidyl peptidase |
| **S2224** | UK 383367 | Procollagen C Proteinase |
| **S7157** | Ilomastat (GM6001, Galardin) | MMP |
| **P3183** | WAY-659100 | hormone sensitive lipase (HSL); metalloprotease |
| **P2039** | WAY-110969 | tyrosinase |
| **S2268** | Baicalein | P450 (e.g. CYP17), endopeptidase |
| **S2269** | Baicalin | GABA Receptor, endopeptidase |
| **S7382** | Phosphoramidon Disodium Salt (PDS) | endopeptidase |
| **S7378** | AEBSF HCl | Serine Protease |
| **P2246** | WAY-313299 | JAMM protease |
| **S9159** | Momordin Ic | SENP1 |
| **S1528** | LY2811376 | non-peptidic β-secretase(BACE1) |
| **S2619** | MG-132 | proteasome and calpain |
| **S2180** | Ixazomib (MLN2238) | 20S proteasome |
| **S8904** | AZ1 | ubiquitin-specific protease (USP) 25/28. |
| **S2714** | LY411575 | γ-secretase |
| **P2462** | WAY-326259 | thrombin; factor XIa |

**B**

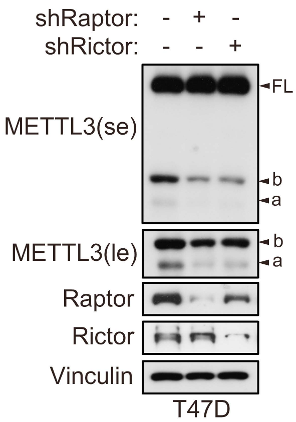


**C**

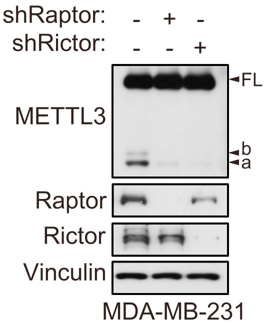
